# Lemborexant Reduces Infarct Volume and Improves Long-Term Functional Recovery in a Murine Model of Ischemic Stroke

**DOI:** 10.64898/2025.12.02.691079

**Authors:** Hee Ra Jung, Shreyas Venkitaraman, Spencer L. Blackwood, Aizad Kamal, Yunhao Jiang, Jake Lee, David Lee, Noor Bibi, Khairunisa M. Ibrahim, Carla M. Yuede, Jin-Moo Lee, Eric C. Landsness

## Abstract

**Introduction:** Recovery following ischemic stroke is highly variable and often incomplete, underscoring the urgent need to develop novel targeted poststroke treatments. While the mechanisms underlying poststroke recovery remain incompletely understood, sleep fragmentation, a common consequence of stroke, has been linked to worse patient outcomes. Lemborexant is a dual orexin receptor antagonist that promotes sleep by suppressing wakefulness and enhancing sleep continuity. We hypothesized that lemborexant would reduce poststroke sleep disturbances and promote recovery in a rodent model of stroke.

**Methods:** We examined the effects of lemborexant (10 mg/kg and 30 mg/kg) and zolpidem (30 mg/kg) on sleep macrostructure, fragmentation, and EEG spectra in both healthy mice and in stroke model mice, which underwent photothrombotic ischemia of the forelimb somatosensory cortex. We also evaluated whether 12 days of drug administration altered infarct volume and functional recovery following the experimental induction of stroke in model mice.

**Results:** Lemborexant treatment (30 mg/kg) increased the percentage of NREM sleep, while preserving sleep continuity, in both healthy and stroke model mice. In contrast, zolpidem increased NREM sleep after stroke, but also increased sleep fragmentation in both groups. Lemborexant treatment at 10 mg/kg and 30 mg/kg significantly reduced infarct volume eight weeks after the induction of stroke. In addition, lemborexant-treated mice showed greater use of the impaired limb four weeks after stroke.

**Interpretation:** These preclinical findings suggest that lemborexant stabilizes sleep and promotes structural and functional recovery following the experimental induction of stroke in model mice, supporting its potential as a novel therapeutic intervention following ischemic stroke.

**Summary for Social Media if Published:**

1. If you and/or a co-author has a X handle that you would like to be tagged, please enter it here. (format: @AUTHORSHANDLE). @EricLandsness
2. What is the current knowledge on the topic? Sleep plays a critical role in structural and functional recovery after stroke, but most pharmacologic sleep aids, such as benzodiazepines and zolpidem, can fragment sleep and impair neuroplasticity. Dual orexin receptor antagonists like lemborexant may offer a newer, mechanistically distinct approach with potential neuroprotective benefits.
3. What question did this study address? This study investigated whether the dual orexin receptor antagonist lemborexant could improve sleep quality, reduce ischemic injury, and enhance functional recovery following stroke in adult mice, compared with the sleep-promoting agent zolpidem.
4. What does this study add to our knowledge? Lemborexant increased NREM sleep without causing fragmentation, reduced infarct volume, and improved motor recovery when administered. These findings suggest that modulating sleep architecture through orexin antagonism during the subacute phase after stroke can promote neural repair and functional recovery.
5. How might this potentially impact on the practice of neurology? Since lemborexant is already FDA-approved for insomnia, these results could be rapidly translated into a therapeutic opportunity to improve stroke recovery through targeted sleep modulation. This approach may shift poststroke care toward integrating neurorestorative, sleep-based interventions during the subacute phase.

## Introduction

Functional recovery following stroke remains highly variable across patients, even when following recommended rehabilitation strategies.^1^ The subacute phase, from three days to six months poststroke, represents an important window for therapeutic intervention that is marked by sleep fragmentation, circadian rhythm disruption, neuroinflammation, and motor and cognitive impairment.^2–7^ Despite this, currently available subacute poststroke treatments have limited ability to enhance endogenous neural repair and remodeling, representing a critical treatment gap.

Sleep disturbances are common after stroke and associated with worse functional and structural outcomes. Clinical studies demonstrate that the presence, severity, and persistence of poststroke sleep disruption predict long-term sensorimotor and cognitive recovery.^8, 9^ Sleep deprivation after focal ischemia results in significantly larger infarct volumes in rodents, as compared to controls with undisturbed sleep.^8, 10^ These findings underscore the critical role of sleep in supporting endogenous poststroke recovery processes.

Despite the important role of sleep in recovery, pharmacological sleep-promoting agents remain underutilized in poststroke care. First-generation sleep-promoting agents, such as benzodiazepines, are positive allosteric modulators of the extensively studied γ-aminobutyric acid type A receptor (GABA_A_R) that enhance GABA binding and chloride influx to induce widespread neuronal inhibition. Although effective for the treatment of anxiety, epilepsy, and insomnia, benzodiazepine treatment is associated with motor and cognitive impairment and amnesic episodes,^11^ so is poorly suited for neurologically vulnerable populations, such as stroke survivors.

Second-generation sleep agents, including zolpidem,^12^ selectively bind to the BZ1 subtype of GABA_A_R, reducing side effects compared to benzodiazepines. Zolpidem treatment during the subacute poststroke phase improved circadian rhythm disruptions, reduced infarct volume, and enhanced functional recovery in rats.^13^ However, zolpidem is associated with side effects that can limit rehabilitation, including next-day drowsiness and psychomotor and memory impairment.^12,14^ Thus, there is a pressing need to develop alternative sleep-promoting therapies that produce fewer side effects.

Orexin peptides A and B regulate wakefulness by binding to orexin receptors 1 and 2 (OX1R and OX2R), representing an alternative target for sleep modulation after stroke.^15–18^ Dual orexin receptor antagonists (DORAs) are FDA-approved for insomnia and improve sleep with fewer side effects than GABA_A_R-binding drugs. DORAs increase NREM sleep and reduce sleep latency,^19–22^ and ongoing clinical trials are assessing their potential to prevent tau accumulation in Alzheimer’s disease.^23^ Lemborexant is the most efficacious DORA for improving subjective total sleep time and reducing wake after sleep onset.^24^ Given the ability of DORAs to enhance sleep with few adverse effects, their benefits in other neurological disorders, and lemborexant’s relative efficacy, we hypothesized that lemborexant would improve sleep and promote functional recovery following stroke.

In this study, we compared the effects of lemborexant and zolpidem on sleep, neurophysiology, and functional recovery in healthy and stroke-affected mice. We found that both drugs selectively increased NREM sleep in healthy mice, but zolpidem increased sleep fragmentation and reduced low-frequency EEG activity. After stroke, lemborexant increased NREM sleep while maintaining continuity, whereas zolpidem again produced fragmentation. Finally, 12 days of poststroke lemborexant treatment reduced infarct volume and improved functional recovery. These findings support orexin antagonism as a therapeutic strategy for stroke recovery.

## Materials and Methods

### Animals

We used male C57BL/6JRj mice aged 12 weeks. Mice were maintained on a 12-hour light/dark cycle with lights on and lights off defined as Zeitgeber Time (ZT) 0 and ZT 12, respectively. All experiments complied with the U.S. Public Health Service Policy on Humane Care and Use of Laboratory Animals and were approved by the Washington University Institutional Animal Care and Use Committee.

### EEG/EMG Surgeries and Stroke Induction

We fabricated custom 6-pin EEG electrode arrays by trimming receptacle connectors (Digi-Key, #ED85100-ND) into 2×3 blocks. We coated the pin shafts with nail polish to provide electrical insulation, cut the male pins to conform to the skull curvature, then applied a conductive silver solution to the exposed ends. A separate 2×2 connector block was prepared for the ground, reference, and EMG channels. The ground and reference pins were processed using the same procedure described above. For the EMG wires, we prepared two 1.5-cm, 32-AWG silver wires (A-M Systems, #790500) by stripping insulation from both ends. We looped one end of each wire and soldered the opposite end to the designated EMG pins at a 45° angle.

We implanted 12-week-old mice with EEG/EMG electrodes under isoflurane anesthesia. Briefly, we placed six 0.46 mm EEG electrodes via burr hole craniotomy, with conductive tips resting on the dura (ML: ±0.65 mm; AP: +1.20 mm, –0.10 mm, –1.40 mm), and secured them with flowable dental composite resin (Tetric N-Flow, Ivoclar Vivadent, Liechtenstein). For EMG implantation, we anchored 2×1 blocks to the skull (ML: ±0.65 mm; AP: –5.50 mm) using the same resin, then subcutaneously tunneled the wires into the cervical and thoracic muscles.

We induced ischemic stroke in mice using optical fiber-induced photothrombosis.^6,7^ Briefly, we implanted a unilateral fiber optic cannula with a ceramic ferrule (RWD, Shenzhen; 0.50 NA, 200 µm diameter) in the left forelimb somatosensory cortex (ML: –2.20 mm; AP: – 0.10 mm; DV: –1.0 mm). To induce stroke, we intraperitoneally injected mice with 0.15 mL Rose Bengal (10 mg/mL), then after five minutes, we delivered a 532 nm, 15 mW solid-state laser through the implanted fiber optic cannula for 15 minutes while mice remained awake and freely behaving. To prevent light scatter, we sealed the connection between the fiber and patch cord with black nail polish. Mice without histologically confirmed infarcts were excluded from analysis.

We administered buprenorphine (30 mg/kg, 0.03 mg/mL) for analgesia after all surgeries, monitored mice for 72 hours, and allowed one week of recovery in their home cages before experiments.

### Experimental Design

#### Experiment #1: Effect of Sleep-Promoting Drugs in Healthy Mice

Seven mice were singly housed in the tethered EEG/EMG recording system for seven days to adapt and train in oral consumption of drug-infused jelly.^25^ Following this, we treated each mouse with 0.50 g of gelatin containing either vehicle (10 mL/kg 0.5% methylcellulose 400), zolpidem (30 mg/kg), or one of two different doses of lemborexant (10 or 30 mg/kg) at ZT 3 on Days 0, 3, 6 and 9, then monitored sleep for 24 hours via EEG/EMG. A zolpidem dose of 30 mg/kg was used as prior work demonstrates that this concentration produces the most robust sedative and NREM-enhancing effects in rodents.^12, 26^ Lemborexant doses of 10 and 30 mg/kg were selected to span a range that produces robust sleep promotion in rodents and to permit assessment of dose dependent effects on sleep and recovery. To control for order effects, we treated each mouse with each treatment in a randomized order, with a 48-hour washout period between treatments during which mice received daily vehicle treatment. ZT 3 is in the early light/inactive phase in mice, corresponding to evening/overnight administration of sleep-promoting drugs in stroke patients and allowing us to maximize recovery sleep. We maintained consistent circadian dosing and drug doses across experiments to support comparison of pharmacodynamic windows, despite differences in administration route.

#### Experiment #2: Effect of Sleep Promoting Drugs after Ischemic Stroke

After seven days of adaptation and training, we induced a photothrombotic stroke in the left forelimb sensorimotor cortex at ZT 3.^27^ At 72 hours poststroke, twenty-eight mice were randomly assigned to receive 0.50 g of gelatin containing vehicle (10 mL/kg 0.5% methylcellulose 400), zolpidem (30 mg/kg), or lemborexant (10 or 30 mg/kg), with seven mice in each treatment group. Sleep was continuously monitored via EEG/EMG from the time of photothrombosis until 24 hours after drug administration, after which the mice were euthanized.

#### Experiment #3: Effect of sleep modulation on stroke recovery

We group-housed sixty-four mice in an enriched environment, with sixteen mice per cage, assigning groups by drug condition to prevent coprophagy-mediated cross-contamination. Before inducing stroke, we performed baseline behavioral testing using cylinder rearing and Y-maze assays at a consistent ZT 3–7, to control for circadian effects. We induced photothrombosis at ZT 3, then from 72 hours poststroke, randomly assigned mice to receive vehicle (10 mL/kg 0.5% methylcellulose 400), zolpidem (30 mg/kg), or lemborexant (10 or 30 mg/kg) via oral gavage daily at ZT 3:00–3:30 for 12 days. As mice in Experiment 3 did not have EEG/EMG implants and could be safely restrained, oral gavage was used instead of gelatin administration to ensure precise and accurate dosing. We repeated behavioral testing before drug treatment on poststroke day 1, and after treatment on days 15, 28, and 56. We euthanized mice and collected brains for histological analysis 24 hours after the final test. Since treatment groups were housed separately to prevent cross-exposure, experimenters were not blinded during photothrombosis or drug administration but remained blinded during behavioral scoring and data analysis.

#### EEG/EMG Data Acquisition

We recorded EEG and EMG signals with a pre-amplifier (RHD2132, Intan Technologies, Los Angeles, CA) and amplifier (Intan Technologies, Los Angeles, CA), sampled at 1000 Hz, and digitized using OpenEphys software.

#### Sleep Architecture Analysis

We downsampled electrophysiological data to 250 Hz, automatically classified five-second epochs as either Wake, REM, or NREM using AccuSleep,^4,5^ then an expert scorer blinded to condition reviewed and manually corrected the classifications. We calculated vigilance state percentage as the time spent in each state divided by the total recording length (three- or 24-hour windows, depending on the analysis). We determined mean bout duration by averaging the length of contiguous episodes of each state. We defined arousals as transitions from NREM or REM to Wake lasting less than 300 seconds, and NREM latency and REM latency as the time from drug administration to the onset of the first NREM and REM bout, respectively. We calculated circadian index of wakefulness (CI) for the 24-hour post-drug administration window using the following formula: CI = (Percent Wake_Dark Period_ – Percent Wake_Light Period_)/Percent Wake_24-Hour_.^13^

#### Spectral Analysis

We analyzed EEG power spectra in three-hour windows. For healthy mice, we examined the first three hours after drug administration. For stroke experiments, we compared baseline recordings (48–51 hours post-stroke) with post-treatment recordings (72–75 hours post-stroke). The signals were resampled to 250 Hz to ensure consistent temporal resolution across recordings. EEG data and EMG data were bandpass filtered between 0.5–50 Hz and 50–100 Hz, respectively. We estimated power spectral density (PSD) across 0.5–30 Hz using short-time Fourier transform with a five-second Hamming window, and plotted them on a logarithmic frequency scale. To summarize spectral shape, we calculated spectral slope as the regression coefficient of the log-log PSD fit and spectral offset as the y-intercept of that fit.^28^

#### Optical Fiber-Induced Photothrombotic Stroke

We induced ischemic stroke in the left somatosensory cortex using optical fiber–induced photothrombosis (OF-PT)^6, 7^. This method enables region-specific, titratable stroke induction in awake, freely moving animals. The procedure was performed as previously described with minor adaptations for fiber-optic delivery.^29^ Mice without histologically confirmed infarcts were excluded from analysis.

#### Behavioral Assessment

Forelimb motor asymmetry was assessed using the cylinder rearing test following established procedures validated for detecting unilateral deficits after forepaw somatosensory cortex stroke. Forelimb contacts in a cylinder were recorded, and an asymmetry index was calculated as the percentage difference between left and right forepaw use. We used a modified Y-maze assay to assess lateralized head-turning behavior rather than spatial working memory, adapting procedures from established Y-maze protocols used in rodent stroke and neurological models.^30^ Mice were placed in a three-arm plexiglass maze and allowed to explore freely while behavior was video-recorded under infrared illumination. Head-turn direction was quantified using automated tracking in ANY-Maze software (Stoelting), and lateralized bias was calculated as the percentage of leftward turns out of total turns. Mice that did not initiate exploration within two minutes were excluded from analysis.

#### Infarct Volume Quantification

We quantified infarct volume to confirm stroke presence, size, and location. Brains were collected, fixed, sectioned, and Nissl-stained using standard histological procedures described in established protocols.^31, 32^ Serial sections spanning the lesion were imaged using a Keyence BZ-X800/810 microscope, infarct boundaries were delineated in ImageJ, and total infarct volume was calculated as the summed infarct area multiplied by section thickness. Mice without confirmed infarcts were excluded from analysis.

#### Secondary Thalamic Injury (STI) Staining

We assessed secondary thalamic injury (STI) using glial fibrillary acidic protein (GFAP) immunohistochemistry performed according to established protocols.^33, 34^ GFAP-labeled sections containing the thalamus were imaged on a Keyence fluorescence microscope, and animals were classified as STI-positive when dense GFAP immunoreactivity was present in the ventral posterolateral nucleus.

#### Statistical Analyses

For Experiment 1 (healthy mice), we compared absolute vigilance state percentages, bout durations, arousals, NREM and REM latency, circadian index and spectral slope and offset across drug conditions using repeated-measures one-way ANOVA after confirmation of normality by Shapiro–Wilk test, followed by Dunnett’s test to compare each treatment with vehicle. If normality was violated, we used the Friedman test. We performed analyses for both the three-hour post-treatment window and the 24-hour recording period.

For Experiment 2 (acute stroke), we calculated intra-mouse changes in vigilance state percentages, bout durations, arousals, NREM and REM latency, CI, and spectral slope and offset from baseline (48–51 hours poststroke) to post-treatment recordings (72–75 hours poststroke).

We compared these differences across groups using one-way ANOVA with Dunnett’s correction after normality testing using the Shapiro-Wilk test. If normality was violated, we used the Kruskal Wallis test instead.

For Experiment 3 (poststroke recovery), we analyzed behavioral performance at baseline and on poststroke Days 1, 15, 28, and 56, and postmortem infarct volume on Day 57. We normalized behavioral asymmetry scores to poststroke Day 1 performance before statistical testing. We compared infarct volume and behavior scores to vehicle using one-way ANOVA with Dunnett’s test for multiple comparisons. We coded binary STI outcomes (STI-positive vs STI-negative) as 1 or 0 and assessed pairwise differences from vehicle using Fisher’s exact test on 2×2 contingency tables.

Sample sizes were based on prior studies using similar stroke and pharmacological paradigms and were sufficient to detect medium-to-large effect sizes in behavioral and histological outcomes (α = 0.05, power = 0.8). No formal power analysis was performed before data collection. To account for multiple comparisons within each experiment, we applied the Benjamini–Hochberg false discovery rate (FDR) procedure with q = 0.05 across related outcome measures separately for each experimental cohort based on independent cohorts, time windows, and hypothesis families. Within each cohort, we ranked p-values from outcome measures and compared them to FDR thresholds. We considered effects with adjusted p-values below threshold significant. We performed all statistical tests in GraphPad Prism (GraphPad, San Diego, CA), and considered results significant at p < 0.05, with FDR correction applied as described above.

### Results Lemborexant increases NREM sleep in healthy mice

In a randomized cross-over design, we treated seven healthy mice with vehicle, zolpidem (30 mg/kg), or lemborexant (10 or 30 mg/kg), and monitored sleep using EEG/EMG for 24 hours, with a 48-hour washout between treatments (**Figure 1A**). Prior studies suggest that DORAs exert their strongest effects within the first 3–6 hours after dosing.^20, 35, 36^ During the three hours post-treatment, lemborexant-(10 and 30 mg/kg) and zolpidem-treated mice (30 mg/kg) produced directionally higher percentage of NREM sleep, relative to vehicle controls, but did not meet significance after correction (p > 0.05; **Figure 1B**). None of the drug treatments significantly altered the percentage of REM sleep during this interval (**Figure 1C**). Mice treated with 30 mg/kg lemborexant showed directionally lower NREM and REM latency relative to vehicle, but these differences did not reach significance (p = 0.086; **Figures 1D–E**).

**Figure 1.**
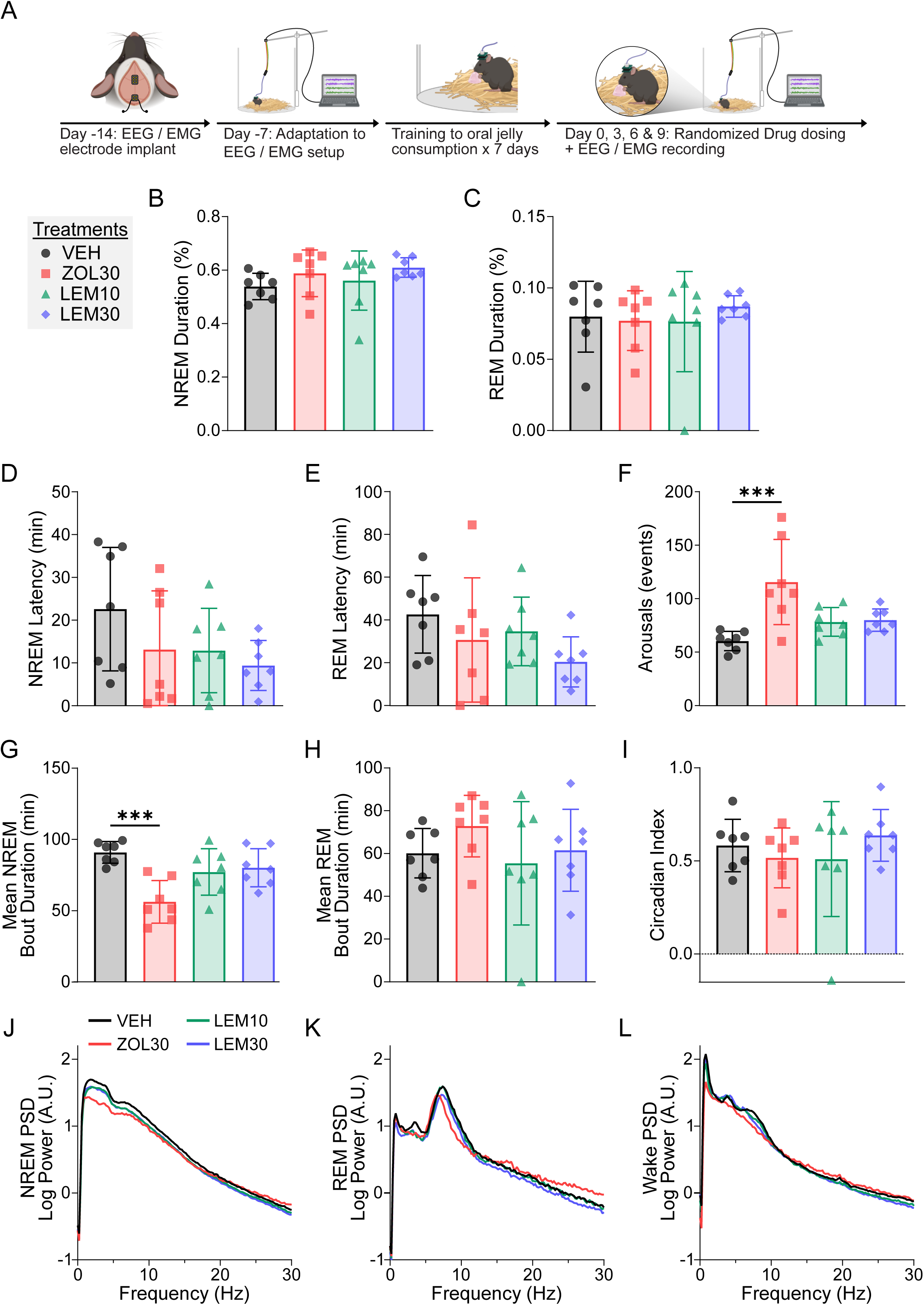
Lemborexant increases NREM sleep and preserves continuity in healthy mice, while zolpidem fragments sleep and reduces NREM EEG power. (A) Experimental timeline: mice underwent EEG/EMG implantation, recovery, and adaptation (Days-14 to 0), then were trained to consume drug-infused jelly. From Day 0, mice received randomized single doses of vehicle, zolpidem (30 mg/kg), or lemborexant (10 or 30 mg/kg) with two-day washout periods between conditions, followed by EEG/EMG recording. (B–C) Percent duration of (B) NREM and (C) REM sleep during the three-hour post-treatment period. (D) NREM and (E) REM latency from treatment to sleep onset. (F) Number of post-treatment arousals, defined as transitions from NREM or REM sleep to wakefulness lasting less than 300 seconds. (G–H) Mean post-treatment (G) NREM and (H) REM bout duration, calculated as the average length of contiguous episodes. (I) Circadian Index. (J–L) Power spectral density (PSD) plots comparing vigilance states across treatments during (J) NREM, (K) REM, and (L) Wake (A.U. = arbitrary units).

When averaged across the 24 hours post-treatment, lemborexant-(10 and 30 mg/kg) and zolpidem-treated mice did not show significant differences in total NREM sleep, relative to vehicle (**Supplementary Figure 1A**). In contrast, zolpidem-treated mice showed significantly lower percentage of REM sleep, relative to vehicle (p = 0.011; **Supplementary Figure 1B**). The post-treatment courses of NREM and REM sleep across 24 hours post-drug administration are shown in **Supplementary Figure 1C and 1D**.

### Zolpidem increases sleep fragmentation in healthy mice

We assessed sleep continuity by quantifying arousals and bout durations of NREM and REM sleep. At three-hours post-administration, zolpidem-treated mice showed significantly more arousal events (p < 0.001; **Figure 1F**) and significantly lower NREM bout duration (p < 0.001; **Figure 1G**), relative to vehicle controls, while lemborexant did not cause significant differences in arousal frequency or NREM bout duration at either dose, relative to controls. No significant differences in REM bout durations (**Figure 1H**) or circadian index of wakefulness (**Figure 1I**) were detected between treatments and vehicle controls. When averaged across the full 24-hour post-treatment period, no significant differences were observed in arousals or bout durations (**Supplementary Figure 2**), although zolpidem showed directionally higher arousal counts that did not reach significance (p = 0.051). In summary, in healthy mice zolpidem induced measurable fragmentation of NREM sleep, whereas lemborexant maintained continuity and showed directional patterns consistent with more stable sleep.

### Zolpidem reduces EEG power during NREM in healthy mice

Prior work shows that zolpidem alters EEG spectral power across frequency bands in NREM sleep,^37^ whereas the effects of lemborexant remain less defined. In the three hours post-treatment, zolpidem-treated mice showed lower low-frequency power during NREM, compared to vehicle (**Figure 1G**). REM and Wake power spectra were unaffected across all treatments (**Figures 1H and 1I**). To further characterize these spectral changes, we quantified the spectral slope and offset. These measures summarize how power decreases with frequency (slope) and the overall power level across frequencies (offset), often interpreted as reflecting the balance of cortical excitation and inhibition.^28, 38^ Zolpidem-treated mice showed significantly lower NREM spectral slope (p = 0.022) and offset (p = 0.002), whereas lemborexant-treated mice (10 and 30 mg/kg) showed no significant differences, relative to vehicle (**Supplementary Figure 3**).

### Lemborexant and zolpidem increase NREM sleep after stroke

We examined the effects of zolpidem and lemborexant treatment at 72 hours poststroke on sleep (**Figure 2A**). We quantified the change in sleep state percentages from baseline (48–51 hours poststroke) to the three-hour post-treatment period (72–75 hours poststroke; **Figure 2B**).

**Figure 2.**
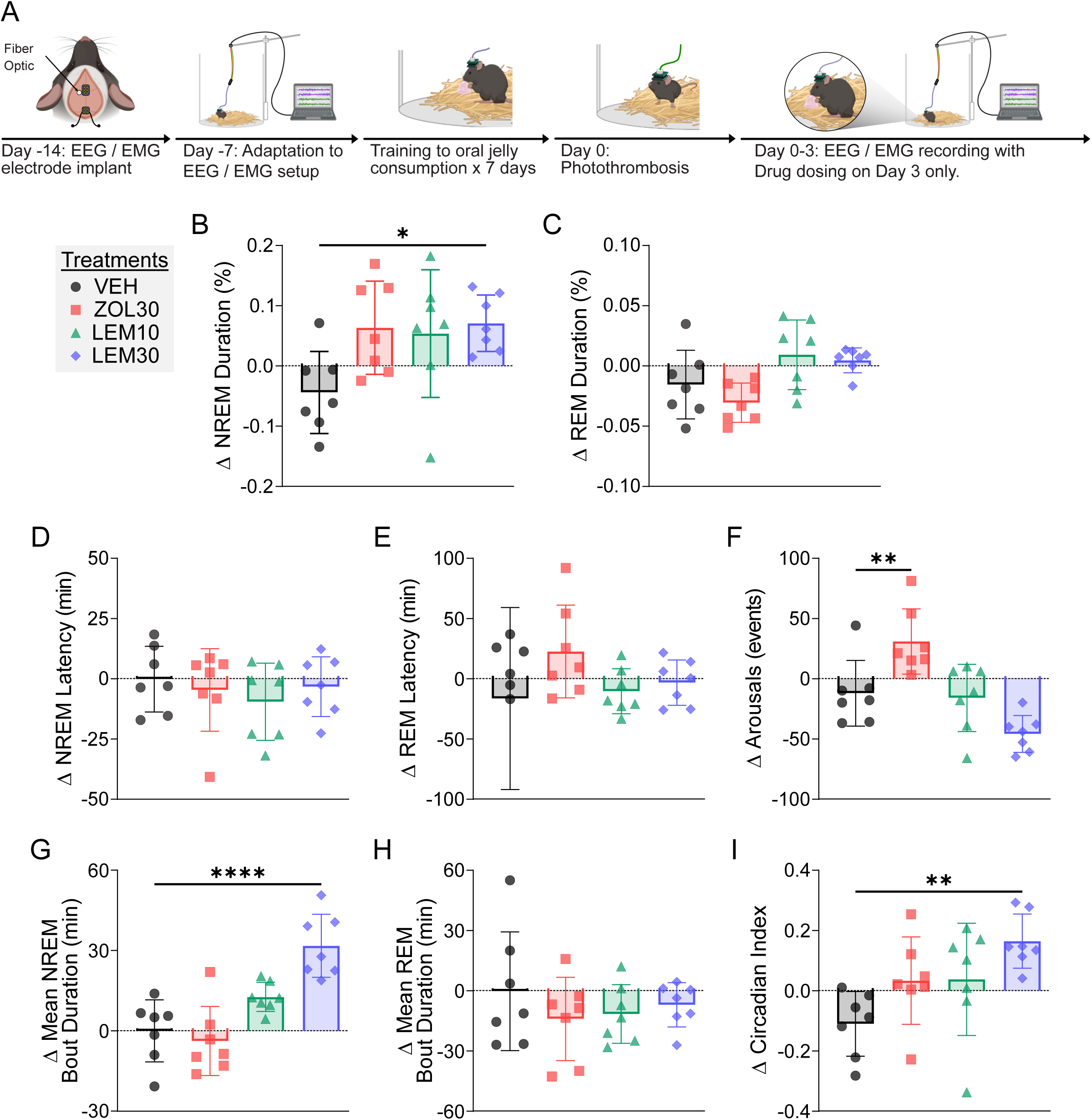
Lemborexant increases NREM sleep and stabilizes continuity after stroke, while zolpidem fragments sleep. (A) Experimental timeline: mice were implanted with EEG/EMG electrodes and a unilateral fiber optic cannula, followed by seven days of recovery and seven days of adaptation with drug-infused jelly. On Day 0, photothrombotic stroke was induced in the left forelimb sensorimotor cortex. On Day 3, mice received a single oral dose of vehicle, zolpidem (30 mg/kg), or lemborexant (10 or 30 mg/kg). Post-treatment changes in EEG/EMG measures were calculated relative to intramouse baseline recordings from hours 48–51 poststroke. (B–C) Percent duration of (B) NREM and (C) REM sleep during the three-hour post-treatment period. (D) NREM and (E) REM latency from treatment to sleep onset. (F) Number of post-treatment arousals, defined as transitions from NREM or REM to wakefulness lasting less than 300 seconds. (G–H) Mean post-treatment (G) NREM and (H) REM bout duration. (I) Circadian Index.

Lemborexant-treated mice (30 mg/kg) showed a significant increase in NREM duration from baseline, as compared to vehicle (p = 0.028). Both zolpidem and 10 mg/kg lemborexant showed directionally higher NREM duration relative to baseline (unadjusted p = 0.042 and p = 0.069, respectively), although neither effect remained significant after correction. No treatments significantly altered the change in percentage of REM sleep (**Figure 2C**), NREM latency (**Figure 2D**), or REM latency (**Figure 2E**) from baseline relative to vehicle during this period. To confirm that these results were not dependent on assessing changes in sleep relative to pre-treatment baseline, we examined absolute post-treatment values during hours 72–75. This complementary analysis showed similar trends across treatment groups and revealed that lemborexant-treated mice (10 and 30 mg/kg) had significantly shorter post-treatment NREM latency, relative to vehicle (**Supplementary Figure 4**).

When averaged across the 24 hours post-administration, no treatment significantly altered the change in NREM or REM duration from baseline, relative to vehicle (**Supplementary Figure 5A and 5B**). The 24-hour NREM and REM time courses (**Supplementary Figure 5C and 5D**) show that both lemborexant (30 mg/kg) and zolpidem induced their most pronounced effects during the 0-6 hours post-treatment period. In summary, lemborexant significantly increased NREM sleep in the early post-treatment period, while zolpidem followed a directionally similar pattern. These early effects were transient and did not persist across the 24-hour period.

### Lemborexant stabilizes sleep continuity after stroke, while zolpidem increases fragmentation

We examined the effects of zolpidem and lemborexant treatment at 72 hours poststroke on sleep continuity by quantifying changes in arousal events and mean REM and NREM bout durations from baseline (48–71 hours poststroke) to the three-hour post-treatment period (72–75 hours poststroke). Zolpidem-treated mice showed a significant increase in arousal events from the baseline 48-71 hour period, compared to vehicle (p = 0.010; **Figure 2F**). Lemborexant-treated mice (30 mg/kg) showed a directionally lower number of arousal events from the baseline 48-71 hour period, compared to vehicle (p = 0.048), though this effect did not survive FDR correction. Lemborexant-treated mice (30 mg/kg) showed a significant increase in mean NREM bout duration from baseline, compared to vehicle (p < 0.001; **Figure 2G**). No treatment significantly altered change in mean REM bout duration from baseline, relative to vehicle (**Figure 2H**). A complementary analysis of absolute post-treatment values during hours 72–75 showed similar trends, with significantly longer mean NREM bout durations in lemborexant-treated mice (10 and 30 mg/kg) relative to vehicle (p = 0.0049 and p = 0.0021, respectively; **Supplementary Figure 6**), while REM bout duration remained comparable across treatments.

When averaged across the 24-hour post-treatment period, zolpidem did not induce a significant difference in change in arousal events from baseline, whereas lemborexant (30 mg/kg) induced a significant decrease in arousal events from baseline, compared to vehicle (p = 0.0010; **Supplementary Figure 7A**). Zolpidem did not induce a significant difference in change in mean NREM bout duration, but lemborexant (30 mg/kg) induced a significant increase in change in mean NREM bout duration from baseline, compared to vehicle (p < 0.001; **Supplementary Figure 7B**). No treatment significantly altered change in mean REM bout duration from baseline, relative to vehicle (**Supplementary Figure 7C**). Finally, we calculated circadian index of wakefulness to assess the effects of each treatment on rhythmic stability after stroke. Lemborexant-treated mice (30 mg/kg) showed significantly increased circadian index from baseline, compared with vehicle (p = 0.003), whereas vehicle, zolpidem, and lemborexant (10 mg/kg) treatment induced no significant differences in change in circadian index (**Figure 2I**). A complementary analysis of absolute post-treatment values during hours 72–75 revealed a similar pattern, with significantly higher circadian index values in 30 mg/kg zolpidem- and 30 mg/kg lemborexant-treated mice (p = 0.0278 and p = 0.0049 respectively; **Supplementary Figure 8**) and 10 mg/kg lemborexant showing directionally higher values. Taken together, these results show that lemborexant maintained more consolidated NREM sleep after stroke, whereas zolpidem was linked to greater sleep fragmentation.

### Lemborexant and Zolpidem Show Limited Effects on EEG Power After Stroke

We examined the effects of zolpidem and lemborexant treatment at 72 hours poststroke on EEG power spectra by quantifying changes in spectra recordings from baseline (48–51 hours poststroke) to the three-hour post-treatment period (72–75 hours poststroke). No treatment significantly altered changes in NREM, REM, or Wake power spectra from baseline relative to vehicle (**Supplementary Figure 9**). Quantitative analysis of spectral slope and offset confirmed no significant effects of zolpidem or lemborexant (30 mg/kg) in any vigilance state. However, lemborexant-treated mice (10 mg/kg) showed significantly higher change in REM spectral slope, compared to control (p = 0.028; **Supplementary Figure 10**).

### Lemborexant Reduces Infarct Volume in the Chronic Phase

Building on the acute effects of these treatments on sleep, we investigated whether repeated administration of zolpidem or lemborexant during the subacute period influenced structural recovery. From three days after stroke, sixty-four mice, with sixteen mice per cage, received daily treatment with vehicle, zolpidem (30 mg/kg), or lemborexant (10 or 30 mg/kg) via oral gavage for 12 days (**Figure 3A**). Both lemborexant doses significantly reduced infarct volume, compared to vehicle (p = 0.001 and p = 0.002, respectively; **Figure 3B**). Zolpidem treatment showed a nonsignificant directional reduction in infarct volume (p = 0.244). These findings suggest that lemborexant treatment in the subacute period after ischemic stroke confers structural neuroprotection.

**Figure 3.**
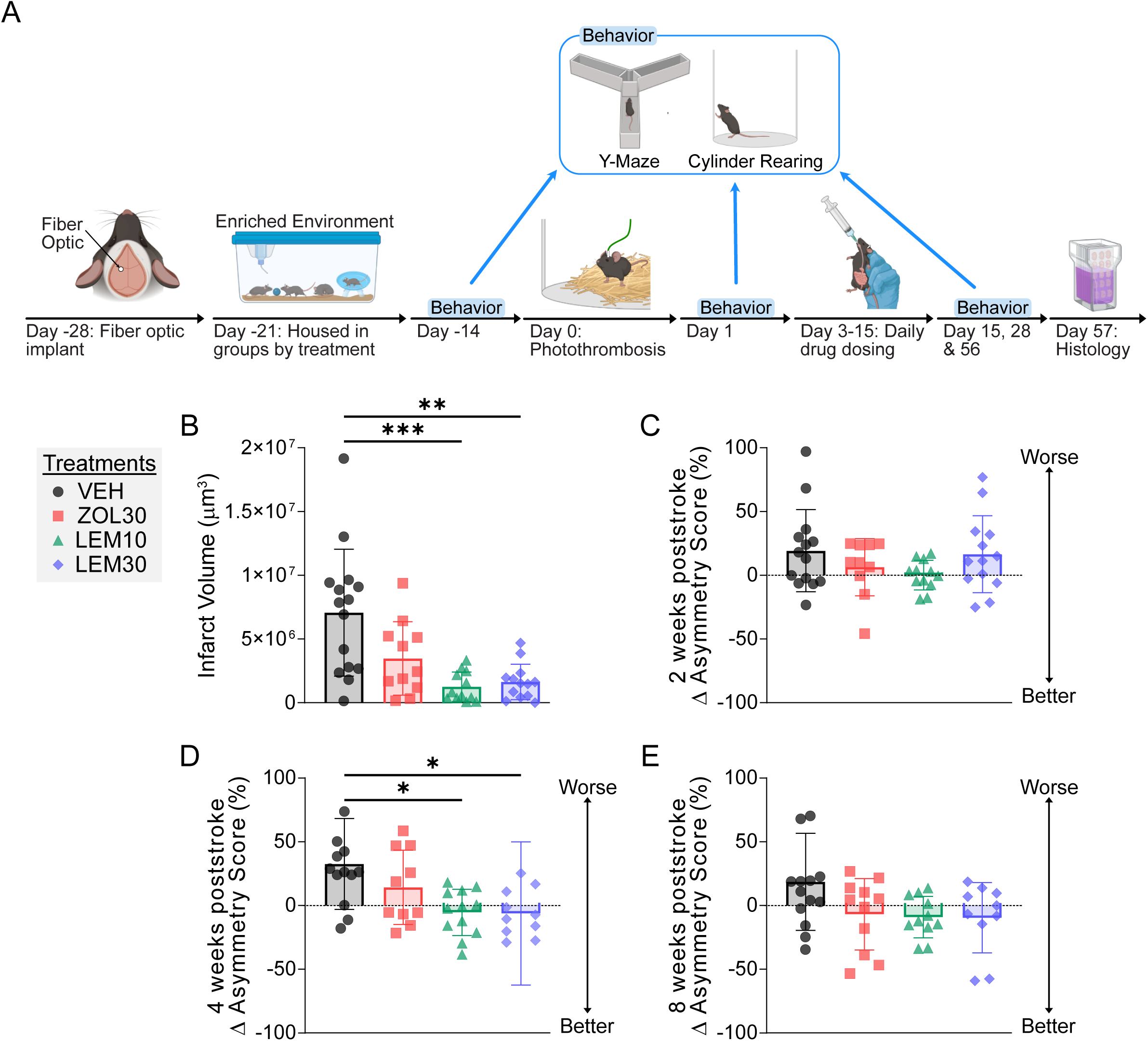
Lemborexant reduces infarct volume and improves forelimb asymmetry after stroke. (A) Experimental timeline: mice were implanted with a unilateral optic fiber for photothrombotic stroke induction on Day −28, followed by seven days of recovery. They were then housed in enriched environments grouped by treatment condition on Day −21. Photothrombosis was induced on Day 0, followed by 12 days of daily oral drug administration beginning 72 hours poststroke. Behavioral assessments were conducted at multiple time points (Day −14, 1, 15, 28, and 56), and tissue was collected for histology on the final day. (B) Infarct volume 30 days after daily oral administration of vehicle (VEH), zolpidem (30 mg/kg; ZOL30), or lemborexant (10 or 30 mg/kg; LEM10, LEM30). Brain sections were stained with cresyl violet, imaged, and quantified to calculate infarct volume. (C–E) Cylinder rearing forelimb asymmetry scores at (C) two weeks, (D) four weeks, and (E) eight weeks poststroke in mice treated with vehicle, zolpidem, or lemborexant. Forelimb asymmetry scores were calculated as (# of frames using left forepaw – # of frames using right forepaw) ÷ total forepaw usage × 100 and expressed as change from baseline asymmetry score at 24 hours poststroke.

As infarct volume does not capture all aspects of injury progression, we also evaluated STI, a marker of remote neurodegeneration.^39^ Zolpidem (p = 0.317) and both doses of lemborexant (p = 0.155 and p = 0.160, respectively)) showed directionally lower prevalence of STI compared with vehicle, although these reductions were not significant (**Supplementary Figure 11**). Overall, lemborexant showed directionally lower STI in a pattern that paralleled its reduced infarct volumes, although these differences were not significant.

### Lemborexant Improves Forelimb Asymmetry After Stroke

We evaluated the effects of repeated administration of zolpidem or lemborexant during the subacute period on sensorimotor recovery by assessing changes in forelimb asymmetry using the cylinder rearing test, a measure sensitive to ischemia in the forelimb somatosensory cortex,^40^ from baseline (24 hours poststroke) to two, four, and eight weeks poststroke. Asymmetry scores were calculated as the percent difference in left versus right forelimb use during rearing.^41^ Asymmetry scores after stroke did not significantly differ between groups (Kruskal–Wallis p = 0.192), indicating comparable pre-treatment behavioral deficits. No significant differences were detected between groups at two weeks or eight weeks poststroke (**Figure 3C and 3E**). At four weeks poststroke, both lemborexant groups (10 and 30 mg/kg) showed significantly lower change in asymmetry from post-stroke day 1, compared to vehicle (p = 0.012 and p = 0.013, respectively; **Figure 3D**). Zolpidem treatment had no significant effects at any timepoint. Taken together, these findings indicate that lemborexant promotes recovery of forelimb asymmetry after stroke, with the strongest effect observed at four weeks poststroke.

As the Y-maze has not been validated as a sensitive measure of recovery after forelimb somatosensory cortex stroke, we tested whether head-turning behavior might provide additional sensitivity. Asymmetry scores were calculated as the change in percentage of leftward head turns relative to total head turns from baseline (24 hours poststroke) to two, four, and eight weeks poststroke. No significant differences were detected between groups at any timepoint (**Supplementary Figure 12**), suggesting that Y-maze head-turning is not a useful marker of recovery in this model.

## Discussion

In this study, we show that lemborexant consolidates sleep and supports recovery after stroke. Lemborexant increased NREM sleep in both healthy and stroke model mice without increasing fragmentation, whereas zolpidem increased arousals in healthy and stroke model mice, shortened NREM bouts in healthy mice, and decreased low-frequency EEG activity in healthy mice.

Across experiments, REM sleep measures did not significantly differ between groups, except for a significant increase in total REM duration observed in mice receiving lemborexant at 30 mg/kg. Lemborexant treatment in the subacute phase reduced infarct volume and improved recovery of forelimb asymmetry, while zolpidem produced no comparable benefit. These findings demonstrate that lemborexant stabilizes poststroke sleep and confers structural and functional protection, supporting orexin antagonism as a promising therapeutic strategy for ischemic stroke.

Several mechanisms could explain the observed benefits of lemborexant following stroke, including enhancement of NREM-dependent plasticity due to reduced sleep fragmentation, stabilization of circadian rhythms, and attenuation of poststroke inflammation. We discuss each of these possibilities below.

Lemborexant may promote recovery by consolidating sleep and enhancing NREM slow-wave activity. Stroke induces hyperexcitability in peri-infarct circuits, and NREM slow-wave activity guides axonal sprouting, supports neuroplasticity, and predicts recovery.^41–43^ Experimental manipulations that boost slow-wave activity, including optogenetic stimulation or pharmacologic activation of GABAergic systems, promote axonal sprouting, neurogenesis, and recovery.^44–46^ Other agents, including baclofen, melatonin, DORA-22, and S44819, support perilesional neurogenesis, plasticity, and synaptic remodeling in preclinical models.^45, 47, 48^ Consistent with these findings, lemborexant increased NREM bout duration without increasing arousals, suggesting that it enhances recovery by stabilizing sleep.

Lemborexant may also promote recovery by stabilizing circadian rhythms. Stroke disrupts the suprachiasmatic nucleus (SCN), the brain’s central pacemaker that regulates immune and inflammatory pathways.^49^ Loss of SCN activity increases neuroinflammatory responses, whereas restoration of rhythmicity can limit secondary injury. Pharmacological interventions that re-engage circadian signaling improve poststroke recovery,^5, 13^ and agents such as zolpidem can restore SCN c-Fos expression in preclinical models, underscoring the importance of central clock mechanisms.^13^ In our study, lemborexant significantly increased the circadian index of wakefulness after stroke, indicating enhanced rhythmic stability. By consolidating sleep and reducing arousals, lemborexant may normalize SCN output, modulate immune responses, and promote neurorepair.

Lemborexant may also promote recovery by modulating poststroke inflammation. The orexin system regulates immune activity, and loss of regulation worsens inflammation and poststroke outcomes.^50, 51^ There is evidence to suggest that activating orexin pathways reduces proinflammatory cytokine levels, such as IL-1β and TNF-α, limits glial activation, and prevents STI.^52, 53^ However, in a recent study based on a tauopathy model, lemborexant reduced microgliosis and protected against neurodegeneration, supporting a role for orexin antagonism in dampening pathological neuroinflammation.^54^ Although some studies report beneficial effects of orexin activation through distinct pathways,^52^ our results indicate that orexin antagonism after ischemic injury may promote recovery by attenuating excessive inflammatory signaling.

Together, these three mechanisms—enhanced NREM-dependent plasticity, stabilization of circadian rhythms, and modulation of poststroke inflammation—offer a framework for how lemborexant may promote recovery after stroke. While our results do not establish causality, they highlight convergent pathways through which sleep stabilization influences neural repair. Future studies should directly test these mechanisms to define their relative contributions and clarify how to best harness orexin antagonism for recovery.

These mechanistic insights gain additional significance because lemborexant is already available as an FDA-approved treatment for insomnia. Unlike GABAergic agents which increase sleep at the cost of fragmentation and cognitive side effects, such as benzodiazepines and zolpidem, lemborexant stabilized NREM sleep without disrupting continuity, indicating that it may better support neural repair. This favorable profile makes it attractive not only for stroke, but also for other conditions where sleep disruption and neuroinflammation worsen outcomes, including traumatic brain injury and neurodegenerative disease. Optimal dosing strategies and treatment windows need to be defined to support the clinical translation of these findings and to ensure both efficacy and safety in patient populations.

Several limitations should be considered when interpreting these findings. First, while lemborexant did not significantly increase the percentage of NREM sleep in healthy mice, there was a consistent trend toward higher NREM values. Combined with prior studies showing robust sleep-promoting effects of lemborexant,^55^ this pattern suggests our analysis was underpowered rather than contradictory with the drug’s profile. Second, lemborexant improved cylinder rearing asymmetry at four weeks but not at eight weeks poststroke. This pattern likely reflects a ceiling effect in which control animals eventually caught up, whereas lemborexant accelerated recovery. Third, we did not detect drug-related differences in EEG spectral power after stroke, with the exception of a modest REM spectral slope increase at 10 mg/kg lemborexant that requires further investigation. These negative spectral results likely reflect stroke-induced baseline alterations, rather than absence of physiological effect. Additional work will be required to disentangle stroke-related EEG changes from pharmacological modulation. Fourth, we did not directly correlate sleep continuity with infarct volume or behavioral recovery because electrophysiological and long-term recovery experiments were conducted in separate cohorts.

Although this design precludes within-subject mediation analysis, convergent effects across experiments support the central conclusion. Future studies could use non-invasive monitoring, such as PiezoSleep, to enable longitudinal assessment of sleep continuity alongside recovery, but this approach cannot distinguish REM from NREM.^56, 57^ Fifth, while all cohorts received drugs at the same circadian time and dose levels, Experiments 1–2 used voluntary jelly ingestion whereas Experiment 3 used daily oral gavage. These differences in administration route may result in handling stress, which could influence both acute sleep physiology and longer-term outcomes. Sixth, lesion size was not measured prior to treatment, so baseline variability could influence chronic outcomes. However, asymmetry scores did not differ across groups at 24 hours poststroke, suggesting similar initial stroke severity. Finally, our experiments used a photothrombotic stroke model in male mice with a relatively short dosing period. Replication across additional stroke models, in both sexes, larger cohorts, and with extended treatment windows will be needed to confirm generalizability. Despite these limitations, the central finding that lemborexant stabilized sleep, reduced infarct volume, and improved recovery remains robust.

In summary, lemborexant consolidated NREM sleep and improved poststroke outcomes by reducing infarct volume and enhancing functional recovery. These results support the broader concept that targeted modulation of sleep architecture can influence brain repair after ischemic injury. Future studies should replicate these findings across stroke models, clarify the relative contributions of sleep plasticity, circadian regulation, and immune modulation, and determine optimal dosing strategies and treatment windows. Given its established safety profile as an FDA-approved treatment for insomnia, lemborexant is a strong candidate for translation into clinical studies aimed at improving stroke recovery. More broadly, orexin antagonism may hold therapeutic relevance for other conditions in which disrupted sleep and neuroinflammation contribute to poor outcomes, including traumatic brain injury and neurodegenerative disease.

## Acknowledgements

This manuscript was developed with assistance from the Scientific Editing Service of the Institute of Clinical and Translational Sciences at Washington University, supported by grant UL1TR002345 from the National Center for Advancing Translational Sciences. Major support for this work was provided by NIH R01 NS13336501 and K08 NS10929201A1, with additional support from NIH R37 NS110699 and R01 NS120481. Study drug was provided by Eisai Inc. The content is solely the responsibility of the authors and does not necessarily represent the official view of the NIH.

## Author Contributions

### Conception and Design of the Study

H.J., S.V., and E.L. contributed to the conception and design of the study. S.V. developed the three-aim experimental framework, and H.R. refined experimental design for surgical, sleep-monitoring, behavioral, and histological components.

### Acquisition and Analysis of Data

H.J. performed all EEG/EMG surgeries, EEG data collection, development and execution of photothrombosis models (optical ferrule–induced and stereotaxic), behavioral testing, vigilance-state scoring, sleep fragmentation analyses, histological quantification, and behavioral data analysis. S.V. performed jelly dosing, oral gavaging, EEG scoring, spectral and vigilance-state analyses (MATLAB, Python, AccuSleep), behavioral scoring, and assisted with brain slicing. S.B. coordinated EEG system operations, contributed to development of the photothrombosis model, trained S.S. in spectral analysis, and provided Python code for EEG processing and AccuSleep integration. A.K. performed photothrombosis surgeries and assisted with perfusion and brain slicing for infarct quantification. Y.J. processed cylinder rearing video data and contributed to automated scoring refinement. J.L. and D.K. assisted with brain slicing, mounting, staining, and histological preparation; J.L. additionally oversaw secondary thalamic injury quantification. N.B. contributed to jelly dosing, EEG recordings, and provided preliminary piezoelectric sleep data that informed study design. C.Y. contributed preliminary data linking Y-maze behavior to sensorimotor asymmetry and assisted with Y-maze data collection.

### Drafting of the Manuscript or Figures

H.J. and S.V. wrote the manuscript and created the figures. They additionally provided technical input on analysis methods and data interpretation. K.I refined the design of the figures. All authors reviewed and approved the final manuscript.

### Study Group Acknowledgment (non-author contributors)

The authors thank Samira Parhizkaror for assistance with jelly drug preparation and protocol refinement, Ashish Sharma for providing preliminary piezo sleep data and technical support, and Braylen Yuede for frame labeling for the cylinder rearing scoring.

### Data and Code Availability

All data are publicly available at Dataset DOI 10.5061/dryad.95×69p900. All data supporting the findings of this study are also available from the corresponding author upon reasonable request. Sleep stage classification was performed using the publicly available open-source MATLAB toolbox AccuSleep. Custom MATLAB scripts used for EEG spectral analysis and the automated behavioral scoring program developed in-house for cylinder rearing analysis are also available from the authors upon reasonable request.

---

**Supplementary Figure 1.**
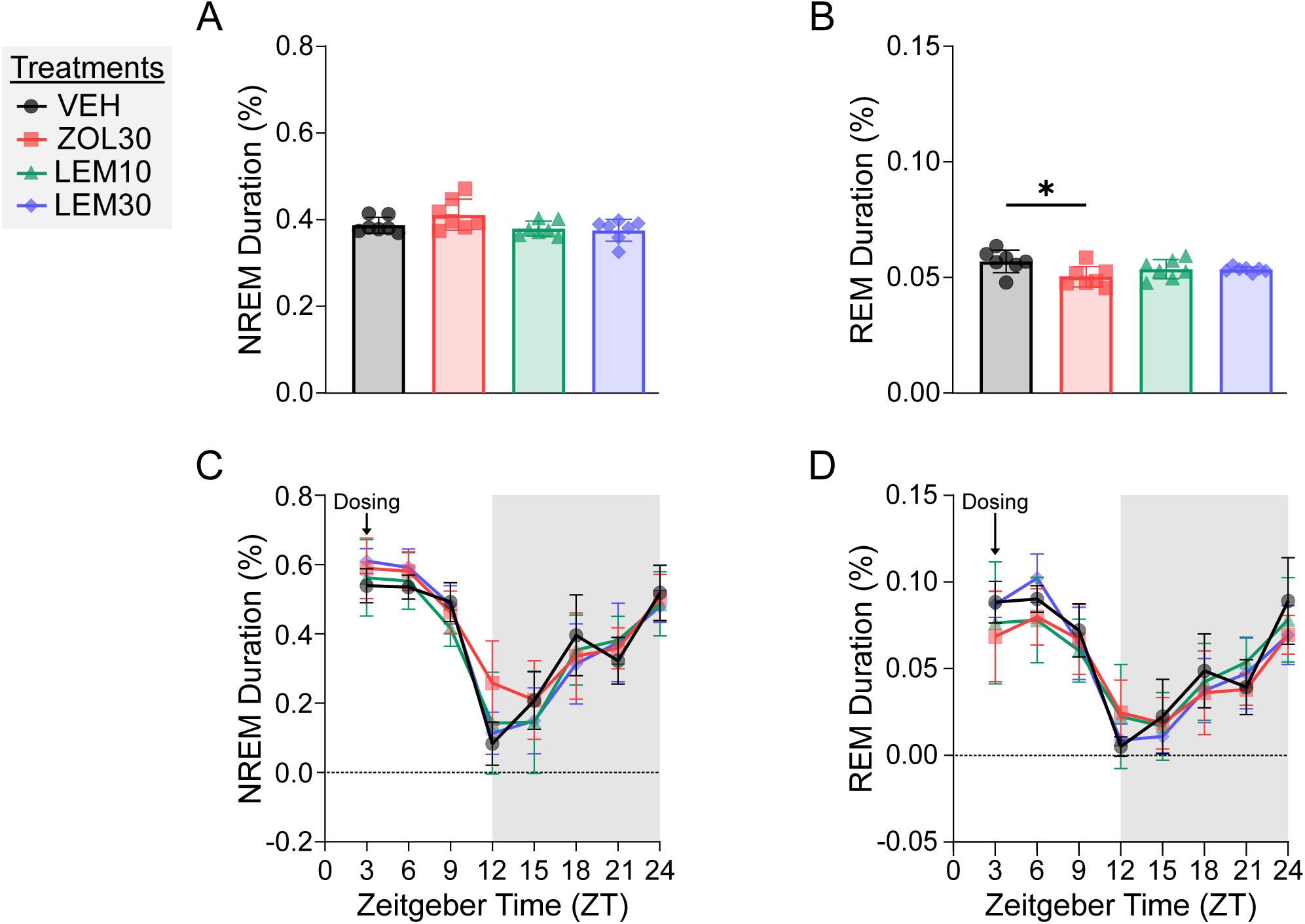
Lemborexant and zolpidem directionally increase NREM sleep in healthy mice without altering 24-hour totals. (A–B) Percent duration of (A) NREM and (B) REM sleep across the 24-hour post-treatment period. (C–D) Time course of (C) NREM and (D) REM sleep plotted in three-hour intervals (ZT0–24).

**Supplementary Figure 2.**
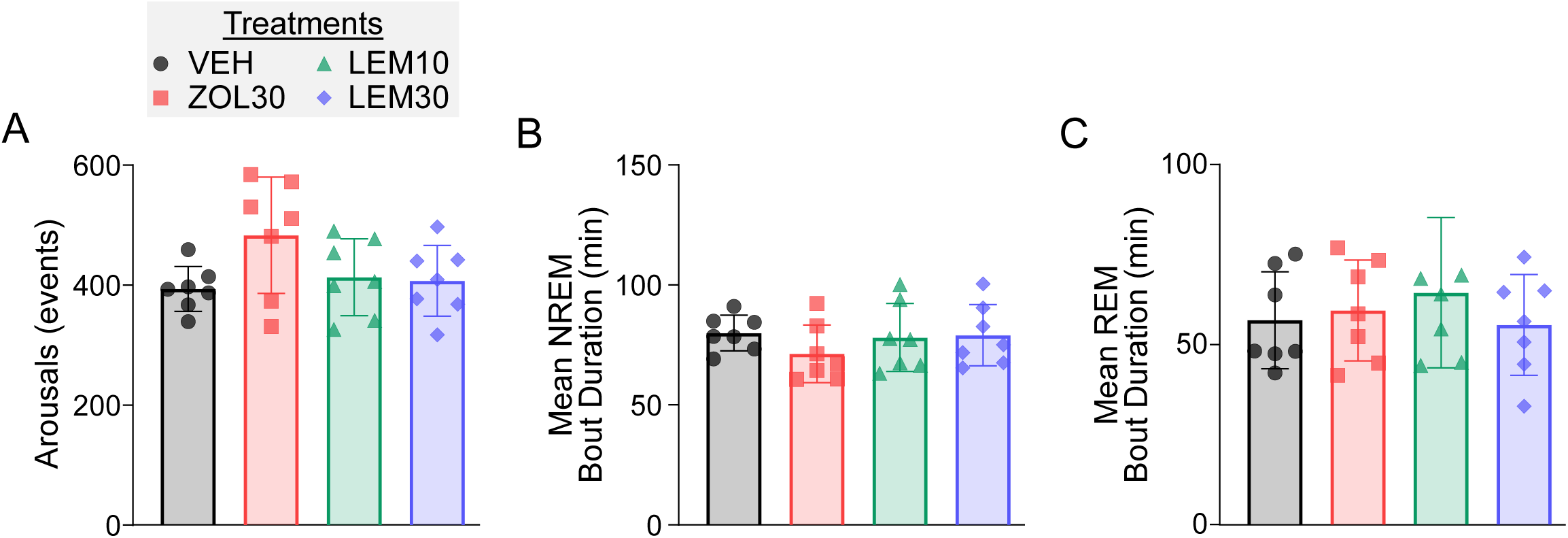
Zolpidem trends toward increased arousals over 24 hours post-treatment, whereas lemborexant maintains sleep continuity in healthy mice. (A) Number of arousals during the 24-hour post-treatment period, defined as transitions from NREM or REM sleep to wakefulness lasting less than 300 seconds. (B–C) Mean bout duration of (B) NREM and (C) REM sleep during the same period.

**Supplementary Figure 3:**
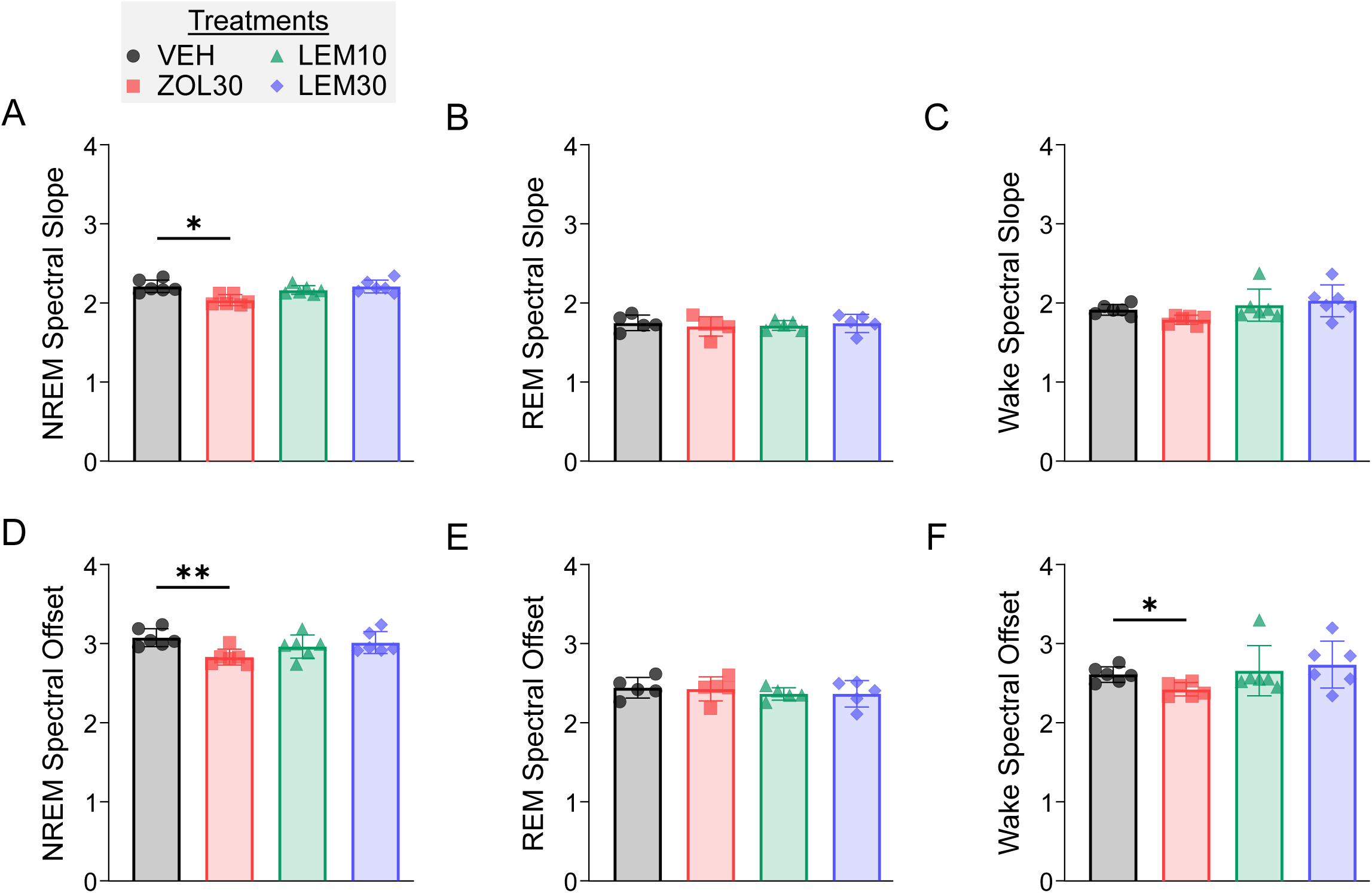
Zolpidem reduces NREM spectral slope and offset, while lemborexant does not alter EEG spectra in healthy mice. (A–C) Spectral slope calculated from the normalized linear fit of the power spectral density (PSD) during (A) NREM, (B) REM, and (C) Wake. (D–F) Spectral offset derived from the y-intercept of the normalized PSD fit during (D) NREM, (E) REM, and (F) Wake.

**Supplementary Figure 4.**
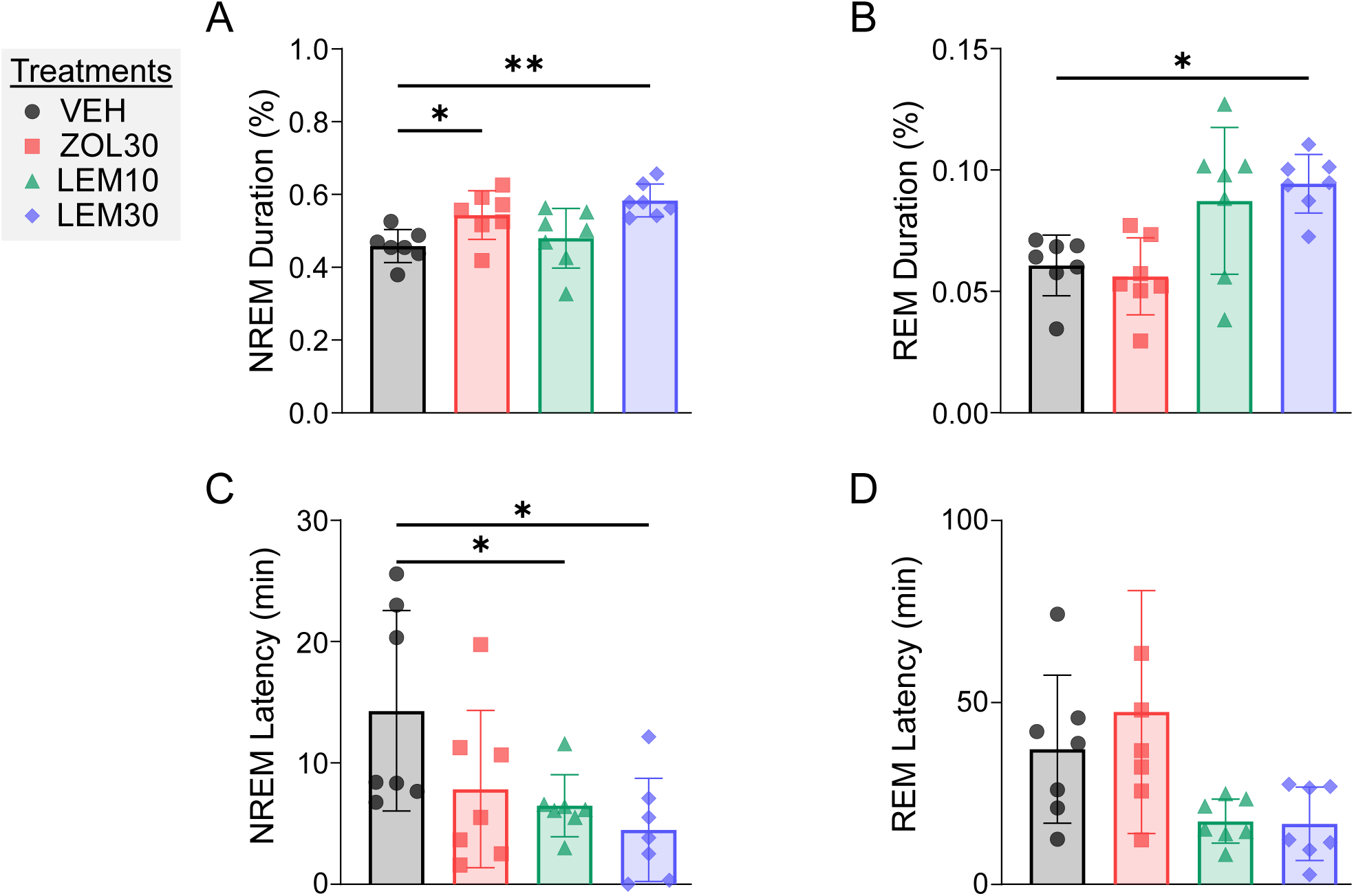
Lemborexant increases NREM sleep and decreases NREM latency after stroke. (A–B) Percent duration of (A) NREM and (B) REM sleep during the first three hours post-treatment (C) NREM and (D) REM latency from treatment to sleep onset.

**Supplementary Figure 5.**
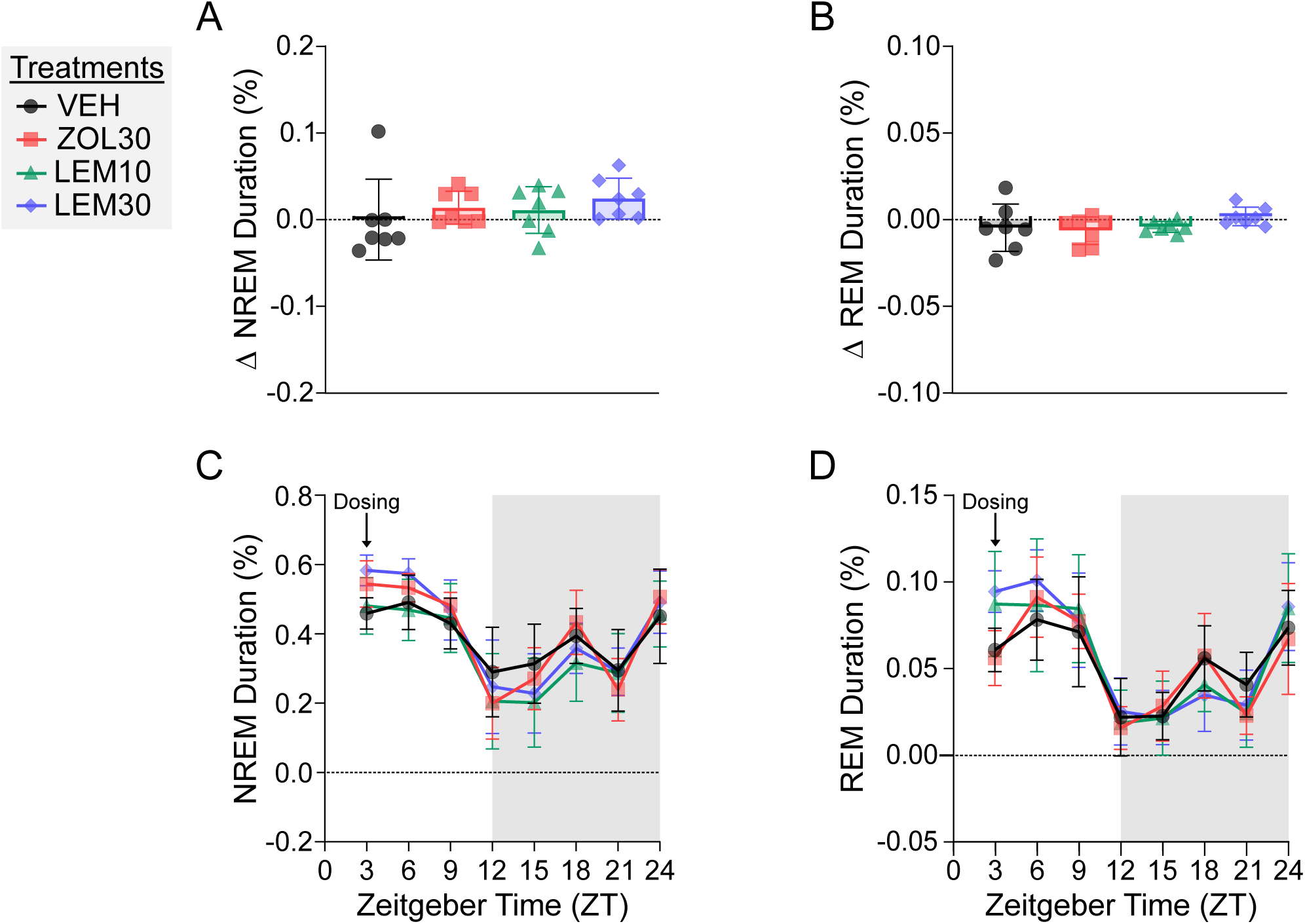
Lemborexant and zolpidem directionally increase NREM sleep after stroke without altering 24-hour totals. (A–B) Relative change in percent duration of (A) NREM and (B) REM sleep across the 24-hour post-treatment period, compared to baseline recordings from hours 48–71 poststroke. (C–D) Time course of (C) NREM and (D) REM sleep across the 24 hours post-treatment period, plotted in three-hour intervals (ZT0–24).

**Supplementary Figure 6.**
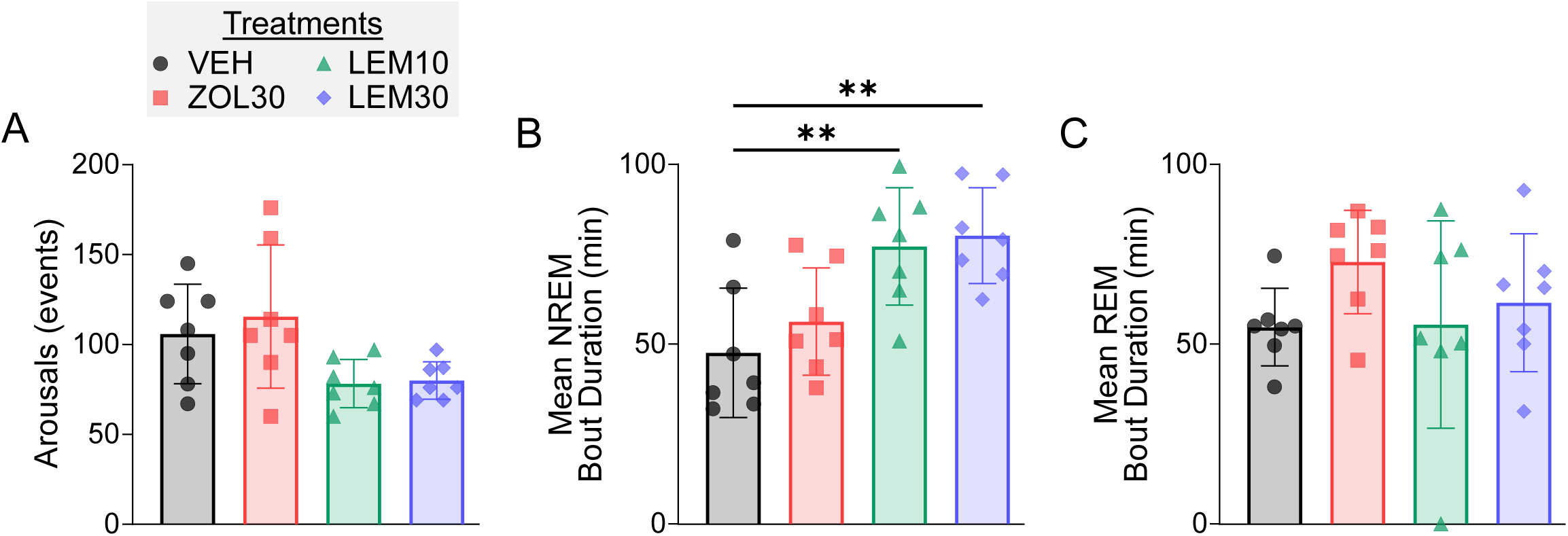
Lemborexant increases NREM duration after stroke. (A) Absolute number of post-treatment arousals during hours 72–75, defined as transitions from NREM or REM to wakefulness lasting less than 300 seconds. (B-C) Mean bout duration of (B) NREM and (C) REM sleep during hours 72–75.

**Supplementary Figure 7.**
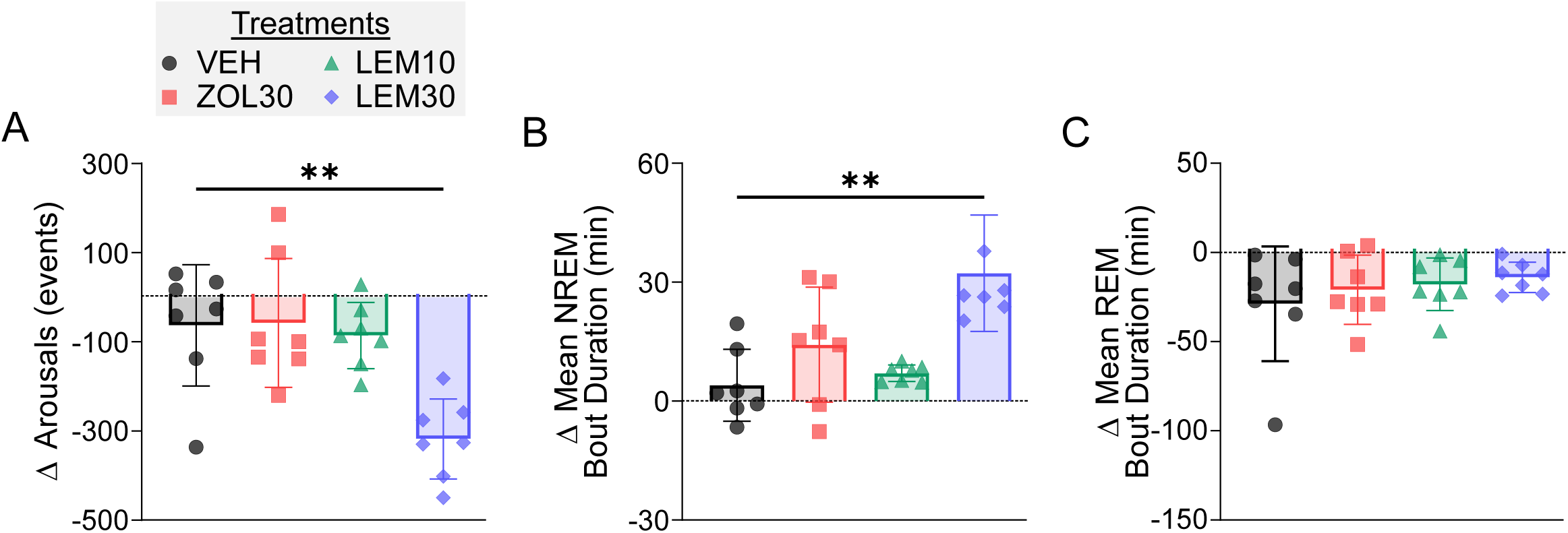
Lemborexant reduces sleep fragmentation over the 24-hour poststroke period, while zolpidem does not. (A) Relative change in the number of arousals from baseline (48–51 poststroke) to the 24-hour post-treatment period, defined as transitions from NREM or REM sleep to wakefulness lasting less than 300 seconds. (B–C) Relative change in mean bout duration of (B) NREM and (C) REM sleep during the same period, compared to baseline.

**Supplementary Figure 8.**
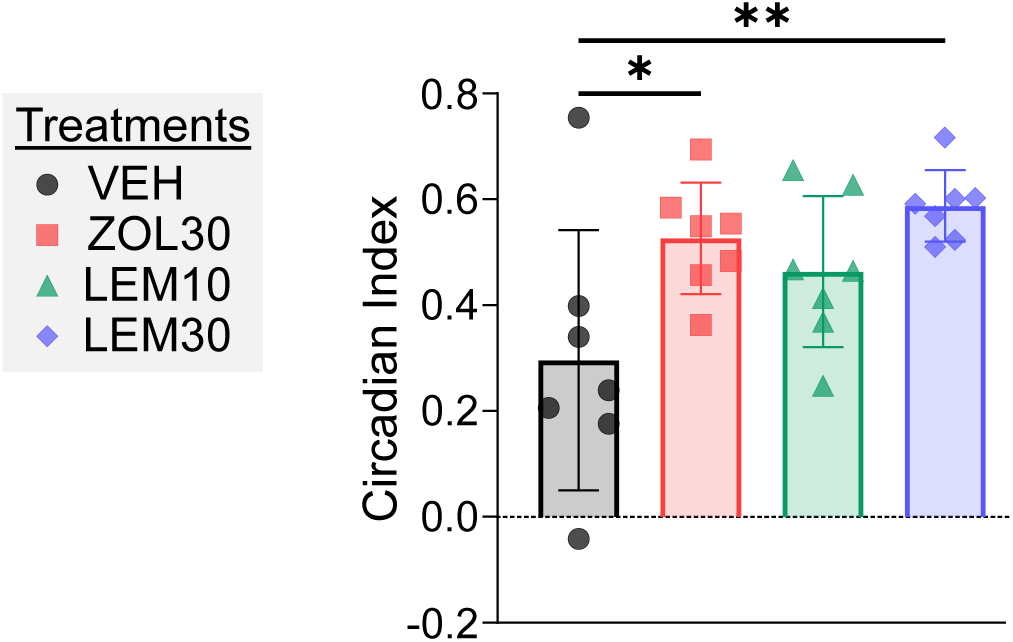
Zolpidem and lemborexant increases circadian index. (A) Absolute circadian index of wakefulness (CI = (Percent Wake_Dark Period_ – Percent Wake_Light Period_)/Percent Wake_24-Hour_) during hours 72-75.

**Supplementary Figure 9.**
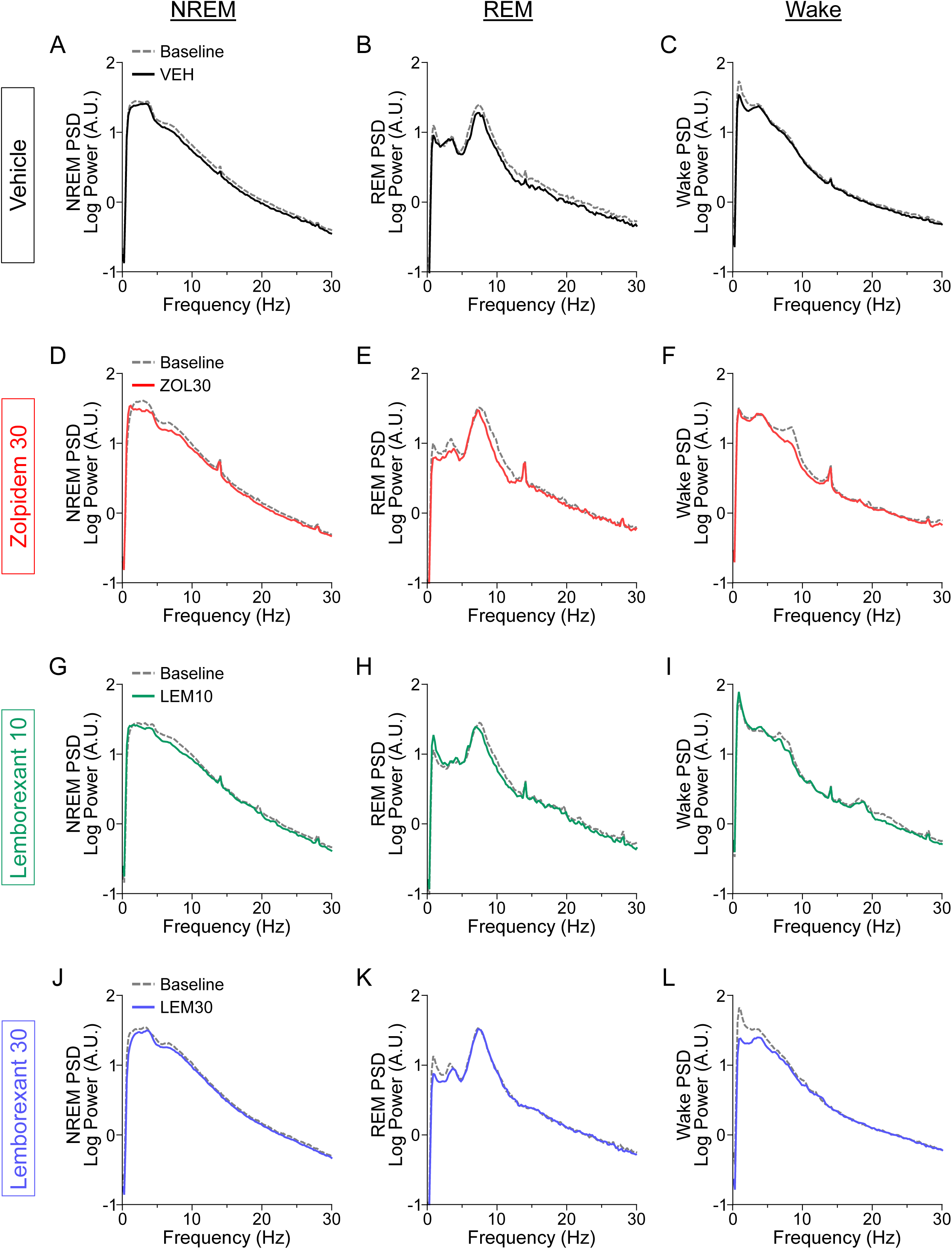
Lemborexant and zolpidem induce no major changes in EEG power spectra after stroke. Power spectral density (PSD) plots comparing baseline recordings (48–51 hours poststroke, gray) with post-drug conditions (72–75 hours poststroke) for each vigilance state and treatment group. Log-transformed PSD was computed across 0.5–30 Hz frequencies. (A–C) Vehicle: (A) NREM, (B) REM, and (C) Wake. (D–F) Zolpidem (30 mg/kg): (D) NREM, (E) REM, and (F) Wake. (G–I) Lemborexant 10 mg/kg: (G) NREM, (H) REM, (I) Wake. (J–L) Lemborexant 30 mg/kg: (J) NREM, (K) REM, (L) Wake.

**Supplementary Figure 10.**
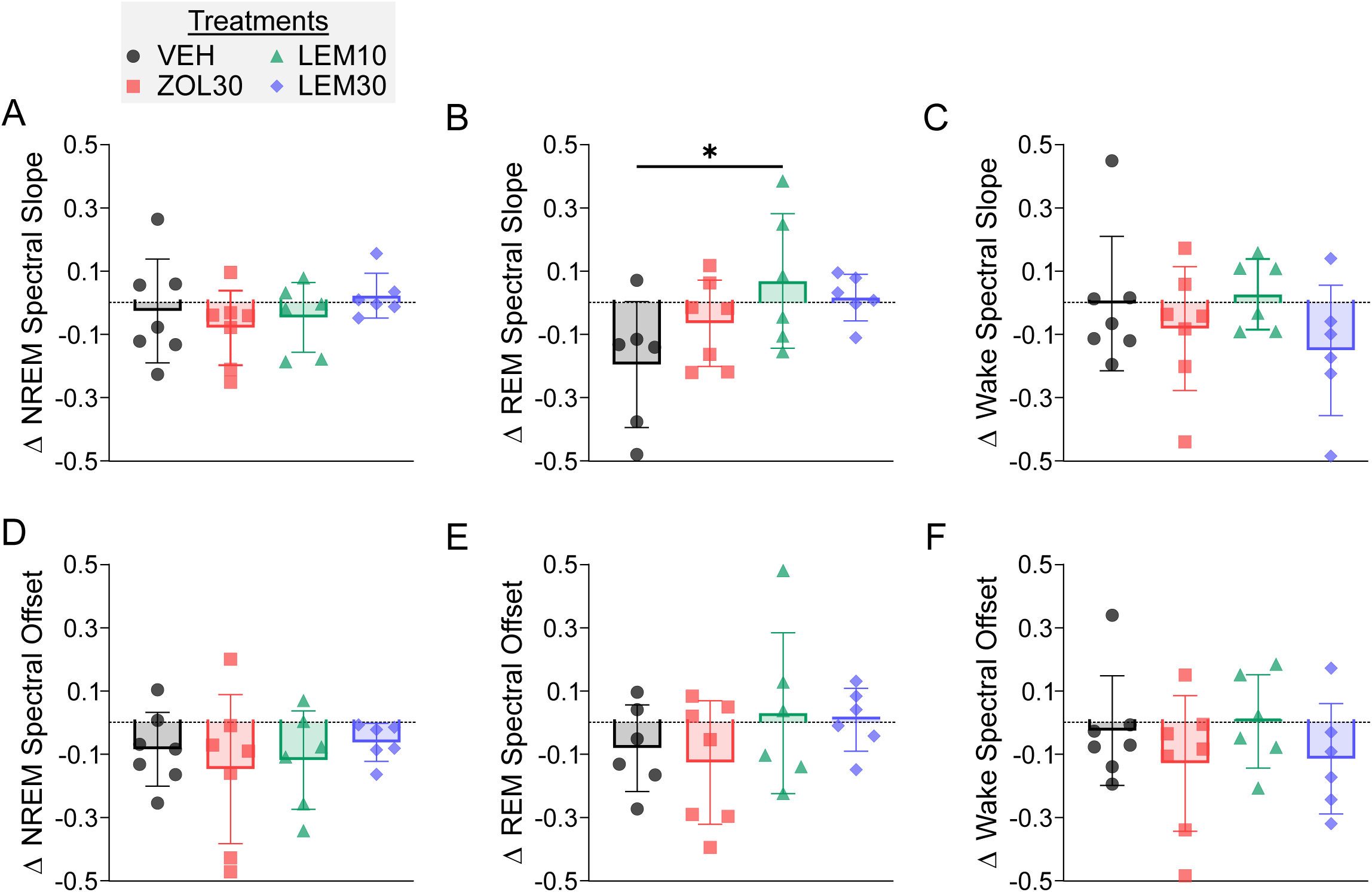
Lemborexant 10 mg/kg increases change in REM spectral slope after stroke, while other drug conditions show no effects. Spectral slope and offset plotted as relative change from baseline recordings (48–51 hours poststroke) to post-treatment (72–75 poststroke hours). (A–C) Relative change in spectral slope during (A) NREM, (B) REM, and (C) Wake. (D–F) Relative change in spectral offset during (D) NREM, (E) REM, and (F) Wake.

**Supplementary Figure 11.**
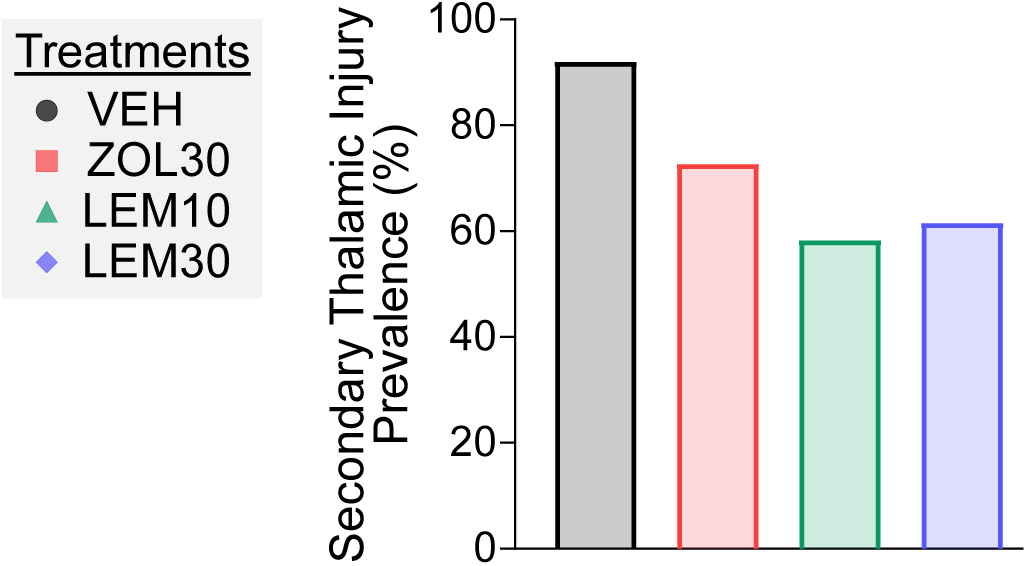
Lemborexant reduces prevalence of secondary thalamic injury after stroke. Prevalence of secondary thalamic injury 30 days after daily oral administration of vehicle (VEH), zolpidem (30 mg/kg; ZOL30), or lemborexant (10 or 30 mg/kg; LEM10, LEM30).

**Supplementary Figure 12.**
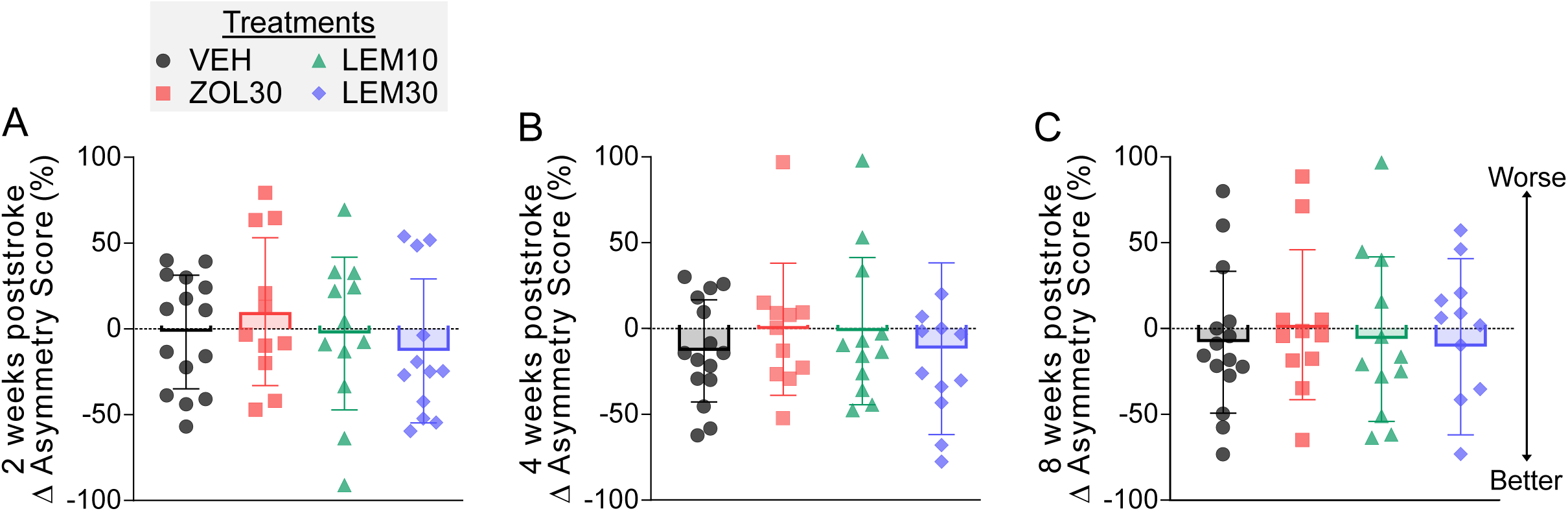
Y-maze head-turning assay shows no treatment-related effects after stroke. Lateralized head-turning behavior in a modified Y-maze at (A) two weeks, (B) four weeks, and (C) eight weeks poststroke in mice treated with vehicle (VEH), zolpidem (30 mg/kg; ZOL30), or lemborexant (10 or 30 mg/kg; LEM10, LEM30). Asymmetry scores were calculated as the percentage of leftward turns relative to total turns, expressed as change from the maximum deficit measured at 24 hours poststroke.

